# Genetic Associations in Four Decades of Multi-Environment Trials Reveal Agronomic Trait Evolution in Common Bean

**DOI:** 10.1101/734087

**Authors:** Alice H. MacQueen, Jeffrey W. White, Rian Lee, Juan M. Osorno, Jeremy Schmutz, Phillip N. Miklas, Jim Myers, Phillip E. McClean, Thomas E. Juenger

## Abstract

Multi-environment trials (METs) are widely used to assess the performance of promising crop germplasm. Though seldom designed to elucidate genetic mechanisms, MET datasets are often much larger than could be duplicated for genetic research and, given proper interpretation, may offer valuable insights into the genetics of adaptation across time and space. The Cooperative Dry Bean Nursery (CDBN) is a MET for common bean (*Phaseolus vulgaris*) grown for over 70 years in the United States and Canada, consisting of 20 to 50 entries each year at 10 to 20 locations. The CBDN provides a rich source of phenotypic data across entries, years, and locations that is amenable to genetic analysis. To study stable genetic effects segregating in this MET, we conducted genome-wide association (GWAS) using best linear unbiased predictions (BLUPs) derived across years and locations for 21 CDBN phenotypes and genotypic data (1.2M SNPs) for 327 CDBN genotypes. The value of this approach was confirmed by the discovery of three candidate genes and genomic regions previously identified in balanced GWAS. Multivariate adaptive shrinkage (mash) analysis, which increased our power to detect significant correlated effects, found significant effects for all phenotypes. The first use of mash on an agricultural dataset discovered two genomic regions with pleiotropic effects on multiple phenotypes, likely selected on in pursuit of a crop ideotype. Overall, our results demonstrate that by applying multiple statistical genomic approaches on data mined from MET phenotypic data sets, significant genetic effects that define genomic regions associated with crop improvement can be discovered.

## Introduction

Almost every crop improvement program assesses the performance of promising germplasm and breeding material via multi-environment trials (METs). The phenotypic data produced by these trials are extremely important guides to growers, private seed companies, and public institutions involved in crop improvement, because combining trial data from multiple years and locations increases the probability of identifying genotypes that perform well or show especially desirable traits (Bowman 1998). Many cooperative testing networks conduct METs to enable cooperators and other interested parties to observe performance over a wider range of environments than if they were only tested locally (Annicchiarico 2002). This supports the identification of advanced lines with stable, high performance in multiple production environments. Amongst many others, crop testing networks that conduct METs include the US cooperative regional performance testing program, the University Crop Testing Alliance, and the Cooperative Dry Bean Nursery (CDBN) (Singh 2000).

Longstanding METs such as the CDBN have often focused on breeding for crop ideotypes, in addition to breeding to eliminate defects and to select for yield. Donald (1968) defined a crop ideotype as an idealized plant with trait combinations expected to produce a greater yield quantity or quality. In contrast, approaches that eliminate defects or select for yield do not consider desirable combinations of traits; thus, these approaches only produce desirable combinations by chance. Selection for an ideotype involves selection for correlated traits, and could lead to substantial pleiotropy, where a single gene affects multiple traits. METs like the CDBN that were used to select for specific crop ideotypes could provide insight into the genetics of trait correlations in crop genomes.

Though METs are often used to measure genetic gain over time (Graybosch and Peterson 2010; Vandemark *et al*. 2014), the vast majority of METs are designed to measure phenotypic responses to a broad set of targeted growing environments. The experimental designs of METs can pose substantial analytical challenges to additional, unplanned genetic analyses. METs typically produce sparse data matrices of phenotypes across germplasm entries, locations, and years (Fig. 1). The frequency of different germplasm entries may vary as part of the normal selection process. Thus, entries with good performance are often tested in more locations and years than those with poor performance. With the exception of few standard checks, the set of genotypes tested each year typically varies, with most genotypes tested in only one or two years. In addition, the total number of genotypes tested each year can vary substantially, and this number is typically too small for genome-wide association on any one year’s data alone. Over the years, MET cooperators can also join or leave the network and add or drop MET sites or phenotypes due to changes in research focus, personnel, or funding. All of these variations make METs into large unbalanced datasets that need to be handled properly for genetic work. Genetic analyses of MET germplasm can also be hampered by the difficulty of obtaining and genotyping previously evaluated entries, particularly entries with poor trial performance that were not tested further. This difficulty may bias or prevent studies that require genetic diversity to explain phenotypic variation, such as genome-wide association studies. In contrast, field experiments designed for genetic studies assess complete, balanced designs, and produce data matrices of phenotypes across genotypes and environments with few or no missing cells. Ideally, the number of genotypes is identical across all environments, and a minimum of a few hundred genotypes are tested in each environment. Each genotype is also tested an equivalent number of times across sites and years.

**Figure 1.**
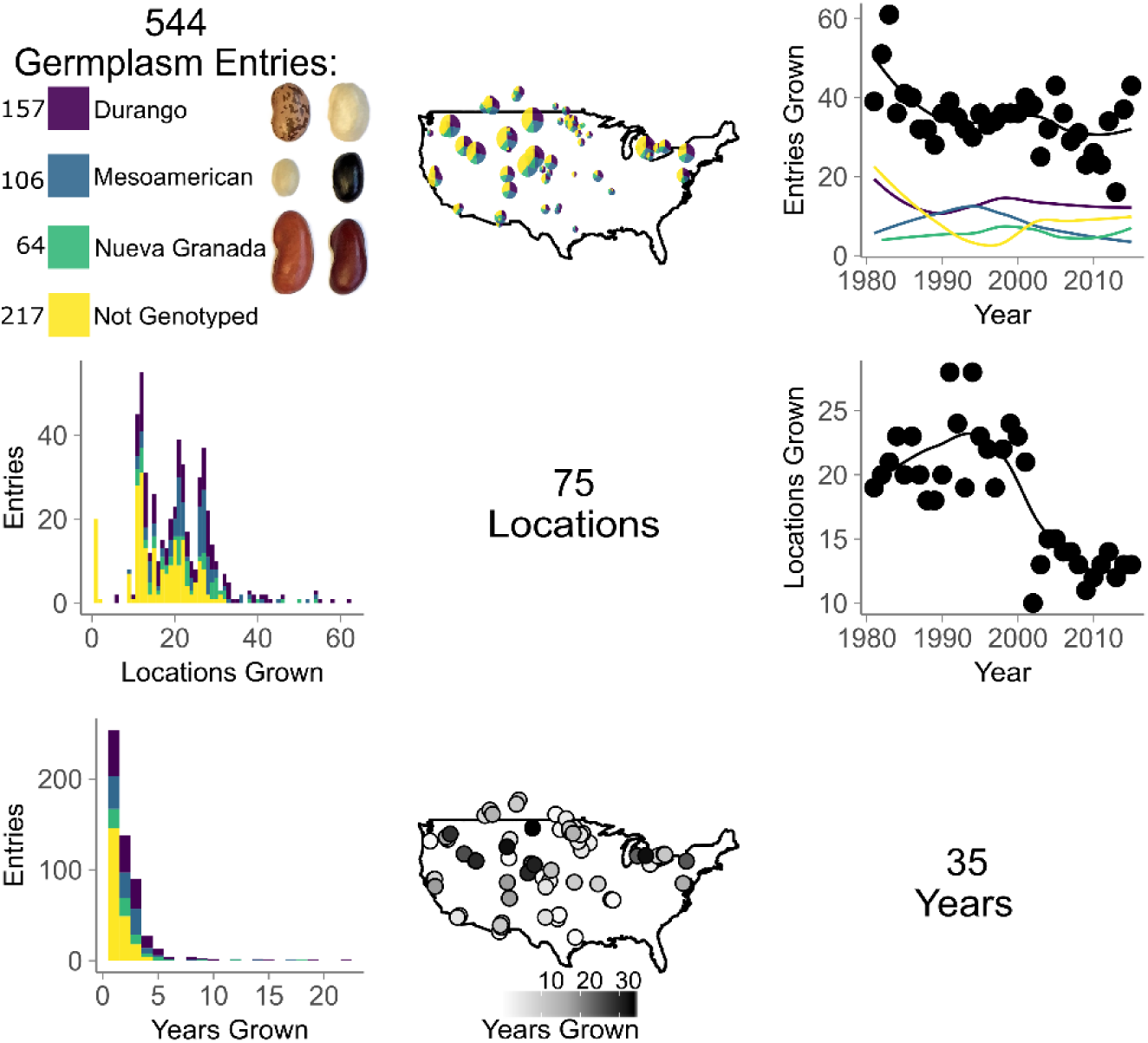
The Cooperative Dry Bean Nursery (CDBN) is an extensive multi-environment trial with hundreds of germplasm entries for common bean (*Phaseolus vulgaris*) grown at 75 locations over 35 years. The relationships between germplasm entries (first row and column), locations (second row and column), and years (third row and column) used in the CDBN are shown in the grid. Purple represents genotyped germplasm entries from the Durango race of the Mesoamerican gene pool, blue represents genotyped entries in the Mesoamerican race of the Mesoamerican gene pool, green represents genotyped entries in the Nueva Granada race of the Andean gene pool, and yellow represents entries that were not genotyped. Beans of the two most commonly genotyped market classes from each race are pictured to the right of the color key, and the number of entries of each race that were genotyped is displayed to the left of the color key. On the maps of CDBN locations, pie chart size is scaled relative to the total number of entries grown at that location, and circle saturation represents locations with a greater number of years of data available.

Despite these analytical issues, METs often produce decades of phenotypic data, which gives them substantial appeal for use in genetic analyses of phenotypic variation. Genetic analyses of MET datasets have recently been implemented in several crop species (Hamblin *et al*. 2010; Rife *et al*. 2018; Sukumaran *et al*. 2018). Its nutritional and agronomic importance, long history of multi-environment trials (METs), and emerging genomic tools makes common bean an outstanding species in which to assess METs that might support the genetic analysis of phenotypic variation. Common bean is the most consumed plant protein source worldwide and is a particularly important source of protein in the developing world (Faostat 2015). In North America, common bean improvement efforts remain mostly in the public sector, and over the past 70 years, the CDBN has been a major testing platform for these improvement efforts. The CDBN is the largest MET for common bean in the United States and Canada (Myers 1988; Singh 2000) and CDBN cooperators have collected phenotypic data on over 150 traits for hundreds of advanced breeding lines and released cultivars (hereafter entries) of common bean at over 70 locations (Fig. 1), which produced up to 18,000 recorded data points per trait (Fig. 2a). The traits are of economic and/or agronomic importance to bean producers, and include seed yield, growth habit, seed size, phenology, and disease responses, among others (Fig. 2a, S1).

**Figure 2.**
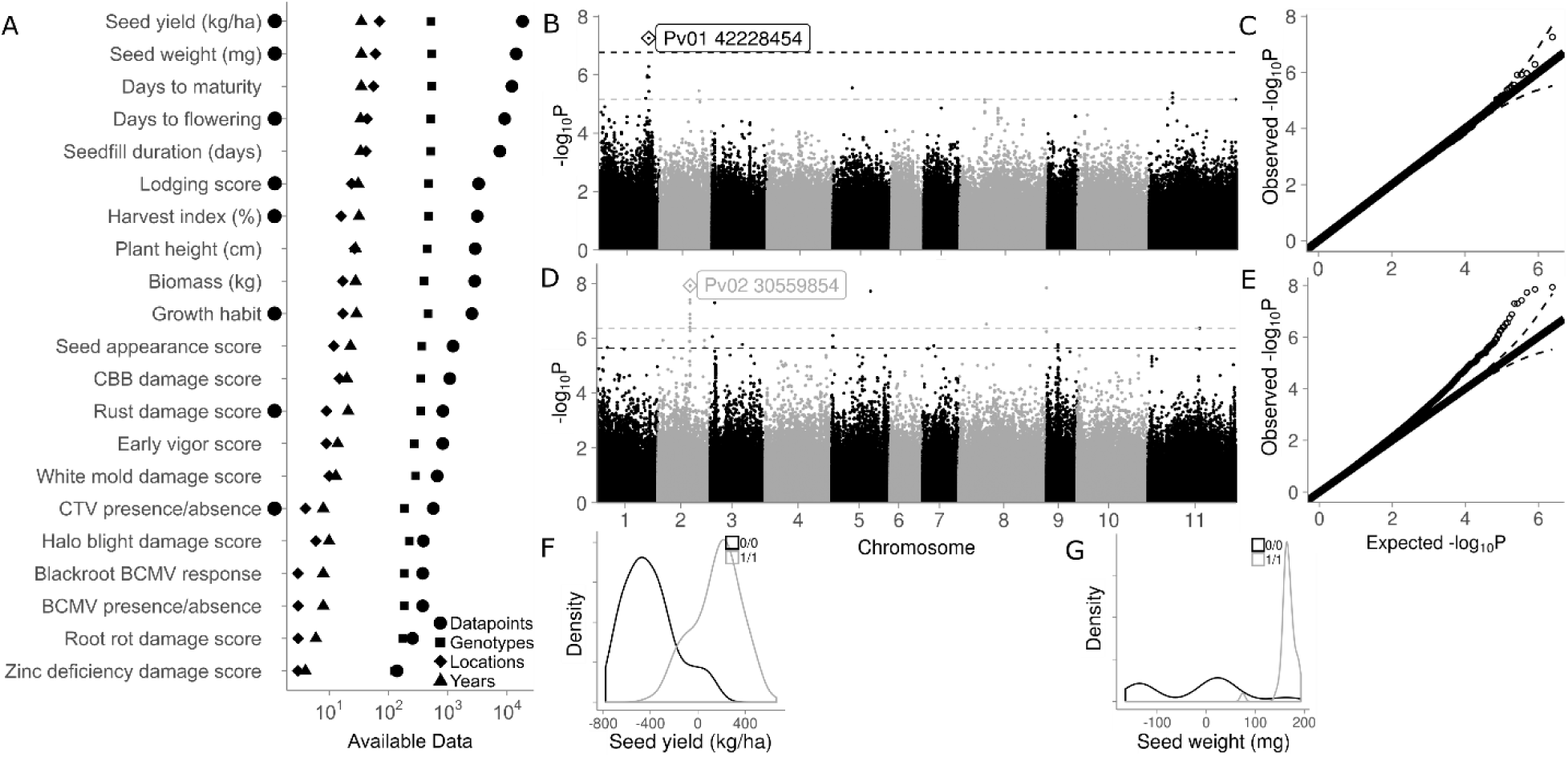
Phenotypic data available in the Cooperative Dry Bean Nursery (CDBN), and genetic variation within the two of these phenotypes with the most data. A) Details of 22 phenotypes present in the CDBN; best linear unbiased predictors (BLUPs) from these phenotypes were used for genome-wide association. Black circles left of the y-axis indicate phenotypes with one or more genetic associations that had *p*-values above the Benjamini-Hochberg false discovery rate. B) Manhattan plot of BLUPs for seed yield (kg/ha) from the CDBN data. The dashed lines are the cutoff values for peak significance. Single-nucleotide polymorphisms above the Benjamini-Hochberg false discovery rate are above the black line, and those in the 0.001 percentile are above the grey line. C) Quantile-quantile (Q-Q) plot of the goodness of fit of the model on seed yield. D) Manhattan plot of BLUPs for seed weight (mg) from the CDBN data, as in B). E) Q-Q plot of the goodness of fit of the model on seed weight. Distribution of the BLUPs for F) seed yield and G) seed weight. Black represents the presence of the reference allele for the top SNP labeled in panel B) and D), respectively, and grey represents the presence of the alternate allele.

More than 500 CDBN entries have been grown since the 1980’s (Fig. 1). These entries include released cultivars and unreleased advanced breeding lines representing most bean types grown in North America. These represent at least thirteen market classes of common bean that group into three major races from two independent domestication events (Mamidi *et al*. 2011) (Fig. 1). Therefore, the CDBN can be used as a representative sample of the genetic diversity being used by North American bean breeders in their programs throughout the last 70 years. However, phenotypic data from the CDBN is sparse and unevenly distributed: the average CDBN entry was grown at only 19 of the 70 locations and in two of the 34 years, with substantial variation in these numbers. CDBN cooperators grew between 16 and 61 of the 500+ entries each year and used ten to 28 of the 70+ locations per year (Fig. 1). Individual CDBN locations grew between eight and 514 entries, with a median of 74 entries. Locations were used in the CDBN for as few as one to as many as 34 years, with a median of five years of participation. Though genotypes are present only intermittently over CDBN locations and years, the vast phenotyping effort on this interrelated set of bean germplasm, when combined with genomic data, offers an excellent opportunity to identify genomic regions affecting phenotypic variation in this species.

Genome-wide association studies (GWAS) have elucidated candidate genes and genomic regions that affect trait variation in many other crop species (Atwell *et al*. 2010; Kirby *et al*. 2010; Mackay *et al*. 2012; Lin *et al*. 2014; Mccouch *et al*. 2016; Macarthur *et al*. 2017; Xiao *et al*. 2017; Togninalli *et al*. 2018) and have recently been implemented in common bean (Cichy *et al*. 2015; Kamfwa *et al*. 2015b; Kamfwa *et al*. 2015a; Moghaddam *et al*. 2016; Soltani *et al*. 2017; Tock *et al*. 2017; Nascimento *et al*. 2018; Soltani *et al*. 2018; Oladzad *et al*. 2019a; Oladzad *et al*. 2019b; Raggi *et al*. 2019). Combining sparse phenotypic data in agricultural datasets to look for pleiotropic effects across conditions has parallels in human biomedical GWAS. In these trials, individual clinics can assess only a subset of human genotypes, and patients are evaluated using institution-specific criteria (Lotta *et al*. 2017; Visscher *et al*. 2017). Human GWAS often look for common variants for common diseases and correct phenotypes for effects of age, sex, and location (Schork *et al*. 2009; Mefford and Witte 2012; Zaitlen *et al*. 2012). Analogously, we seek common, genetically stable variants for important phenotypes evaluated in a MET, corrected for effects of location, year, kinship, and assessment criteria. In human biomedical GWAS, pleiotropic effects of SNPs on multiple diseases have frequently been observed (Sivakumaran *et al*. 2011). Selection for a common bean crop ideotype, with a long hypocotyl, many nodes carrying long pods and without side branches, small leaves, and determinate growth (Adams 1982; Kelly 2001), is known to have led to pleiotropic effects on multiple traits, such as seed yield, biomass, lodging, and plant height (Soltani *et al*. 2016). To study the genetic effects of this aspect of the CDBN selection framework, we used multivariate adaptive shrinkage (mash) to find genomic associations with significant effects on one or more CDBN phenotype (Urbut *et al*. 2019). Mash is a flexible, data-driven method that shares information on patterns of effect size and sign in any dataset where effects can be estimated on a condition-by-condition basis for many conditions (here, phenotypes) across many units (here, SNPs). It first learns patterns of covariance between SNPs and phenotypes from SNPs without strong effects, then combines these data-driven covariances with the original condition-by-condition results to produce improved effect estimates. In this way, mash shares information between conditions to increase the power to detect shared patterns of effects. Mash was originally used for analyses of human biomedical data (Urbut *et al*. 2019) and has yet to be used in an agricultural setting. This analysis method could be used with the rich phenotypic resources of crop METs to understand genetic effects across multiple phenotypes or across multiple locations and years.

Here, we demonstrate that the CDBN MET dataset can be used to make genetic discoveries, despite the sparse nature of the data, by using BLUPs for entries phenotyped in the CDBN. We explore whether this approach can find genomic regions significantly associated with phenotypic variation, and compare associations found with this approach to published GWAS results obtained from more balanced trials. We also explore patterns of genomic associations with significant effects on more than one CDBN phenotype using mash. Our results demonstrate the value of adding a genetic component to datasets such as the CDBN and provide a starting point for future work that explores the genetics of phenotypes evaluated in METs.

## Materials and Methods

### Background principles: processing, digitization and genetic analysis of phenotypic data

MET datasets represent substantial phenotypic resources that can aid in the genetic study of important agronomic phenotypes. Several important steps in preparing the CDBN data for analysis fall under the remit of data science, and specifically involve the data processing steps outlined here. First, when available only from printed reports, the data was rendered machine-readable. Processing of the digitized data next involved cleaning the data to remove inconsistencies and spurious data, then filtering to retain only the relevant data. The data was stored in a consistent form where the semantics of the dataset matched the way it was stored. Then, various data scales for individual traits such as growth habit were standardized to create phenotypes that were more consistent across locations and years. The phenotypic data was next enriched with additional attributes that made subsequent analyses more meaningful, such as germplasm, environment, and crop management information. Then, the data was aggregated to create summary data, by estimating BLUPs for each phenotype. We next used a GWAS modeling approach to determine the genomic regions associated with these data summaries. Finally, we used multivariate adaptive shrinkage (mash) to examine the patterns of overlap between genomic associations with significant effects on one or more phenotype (Urbut *et al*. 2019).

#### Phenotypic data processing

Phenotypic data for entries grown in the CDBN were available mainly as hard-copy reports providing plot averages at named locations. Some reports were available in the National Agricultural Library from the 1950s onwards; however, reports from 1981 onwards had substantial additional available genetic material and were the focus for this analysis (Table S1). Reports from 1981 to 2015 were scanned if not in digital format, digitized using optical image recognition as required, and then reformatted using custom SAS (SAS System, version 9.4, SAS Institute Inc., Cary, NC) scripts that also standardized nomenclature and units of measurement.

Much of the phenotypic data required additional processing to allow comparisons across locations and years. The long timespan and large number of testing locations led to the scoring of 152 traits. Many of these traits represented distinct methods for scoring similar phenotypes; for example, lodging was scored on a percent scale, a 1 to 5 scale, a 0 to 9 scale, and a 1 to 9 scale at different locations and in different years; for this analysis, these lodging traits were standardized to one lodging phenotype on a 1 to 5 scale. From 152 traits reported, 22 phenotypes were standardized for use in GWAS, including eight quantitative phenotypes and fourteen qualitative phenotypes created from visual scores and/or specific measurements (Fig. 2a). The output from the R script used to standardize the phenotypes across locations and years can be found online at http://rpubs.com/alice_macqueen/CDBN_Phenotype_Standardization.

We generated phenotypes associated with location code, year, and genotype information. A total of 70 location codes were created as four-letter abbreviations with the U.S. state or Canadian province abbreviation as the first two letters, and the specific site abbreviation as the second two letters. Five location codes ending in “2” corresponded to a second trial grown at that location and year, usually with a treatment such as drought or disease applied. Location codes were associated with latitude, longitude, elevation, and other location-specific metadata (Table S2), while genotypes were associated with market class and race, as well as the availability of seed from the holdings of CDBN cooperators and single nucleotide polymorphism (SNP) data, where available (Table S2).

In general, location by year (L*Y) combinations with outlier phenotypic values (values above the third quartile or below the first quartile by 1.5 times the interquartile range, or IQR) were removed for every entry in that L*Y combination. Removing outlier L*Y combinations prevented possible bias from linear models using a biased sample of datapoints for a L*Y, while still removing points that, by IQR measures and by knowledge of reasonable ranges for common bean quantitative phenotypes, were likely due to mismeasurement or data entry errors. The specifics of phenotype standardization for all 22 phenotypes are given in the Supplementary Note and the code is available on GitHub at https://github.com/Alice-MacQueen/CDBNgenomics/tree/master/analysis-paper.

### Germplasm: CDBN Diversity Panel and Single Nucleotide Polymorphism Dataset

#### Germplasm recovery and sequencing

To detect genomic regions associated with phenotypic variation in a GWAS framework, it is particularly valuable to have a large amount of heritable phenotypic variation. Thus, it was equally important to include entries from the CDBN with poor seed yields or non-ideal phenotypic traits as high yielding, commercially released varieties. We thus went to considerable effort to obtain seed of unreleased, unarchived materials from the holdings of CDBN cooperators. Germplasm from the entries grown in the CDBN was obtained from multiple sources, including the International Center for Tropical Agriculture (CIAT), the National Plant Germplasm System (NPGS), and three common bean diversity panels, the Mesoamerican Diversity Panel (MDP) (Moghaddam *et al*. 2016), Durango Diversity Panel (DDP) (Soltani *et al*. 2016), and Andean Diversity Panel (ADP) (Cichy *et al*. 2015). Seed was also obtained from holdings of CDBN cooperators, including Mark Brick (Colorado State University), Jim Kelly (Michigan State University), Phil McClean (North Dakota State University), Phil Miklas (USDA-ARS), James Myers (Oregon State University), Juan Osorno (North Dakota State University), and Tom Smith (University of Guelph).

The SNP dataset was created from this germplasm in two ways. First, raw sequence data was obtained from the ADP, DDP, and MDP (Cichy *et al*. 2015; Moghaddam *et al*. 2016) for CDBN entries and all parents of CDBN entries which had been sequenced as part of these panels. The remainder of the CDBN was genotyped using identical methodology to these previous diversity panels, dual-enzyme genotyping-by-sequencing (SCHRÖDER *et al*. 2016). Unfortunately, 39 of the older, unreleased varieties would no longer germinate. For these varieties, we obtained DNA for sequencing by rehydrating sterilized seeds on wetted Whatman paper in petri plates for 2-3 days, then dissecting the embryo from the seed and extracting DNA from the embryo. The DNA from the remaining entries was extracted from young trifoliates. The enzymes *MseI* and *TaqI* were used for digestion following the protocol from Schröder *et al*. (2016). SNPs were called from this raw sequence data using the pipeline found at https://github.com/Alice-MacQueen/SNP-calling-pipeline-GBS-ApeKI. Briefly, cutadapt was used to trim adapters and barcodes (Marcel 2011), sickle adaptive trimming was used to remove ends of reads with quality scores below 20 (Joshi and Fass 2011), bwa mem was used to align reads to V2.0 of the G19833 reference genome found at https://phytozome.jgi.doe.gov/pz/portal.html#!info?alias=Org_Pvulgaris (Li and Durbin 2010; Schmutz *et al*. 2014), and NGSEP was used to call SNPs for the entire set of CDBN entries and all parents in the CDBN pedigrees (Duitama *et al*. 2014). SNPs were imputed using FILLIN in TASSEL. This resulted in the creation of a diversity panel of 327 entries with MET data in the CDBN, (Table S2) with aligned SNP data available on the UT Libraries data repository at doi: <*to be obtained before publication; authors can provide for analysis replication purposes during review>* for use in the CDBNgenomics R package at https://github.com/Alice-MacQueen/CDBNgenomics.

#### Genome-wide association study

To explore consistent genetic effects that could be compared to balanced genetic trials, analyses were performed on genetic BLUPs for each phenotype. BLUPs were calculated in the rrBLUP package in R, using a kinship matrix and treating location and the interaction between location and year as fixed effects. The R code to generate the BLUPs is available on GitHub at https://github.com/Alice-MacQueen/CDBNgenomics/tree/master/analysis-paper. The BLUPs are available in Table S2. For GWAS phenotypes, BLUPs were retained only for CDBN entries phenotyped at least one time in the CDBN. The kinship matrix was calculated using default methods in GAPIT. A total of 1,221,540 SNPs with a minor allele frequency greater than 5% in the CDBN diversity panel were identified and used for the CDBN GWAS. GWAS analyses were performed using compressed mixed linear models (Zhang *et al*. 2010) implemented in GAPIT with the optimum level of compression (Lipka *et al*. 2012). These models used a kinship matrix calculated within GAPIT to control for individual relatedness, and some number of principle components (PCs) to control for population structure. The optimum number of principle components (PCs) to control for population structure was determined using model selection in GAPIT, and by selecting the number of PCs that maximized the Bayesian Information Criterion (BIC). Typically, zero to two PCs were used (Table S3). The final Manhattan plots were created using the ggman R package. Plots of intersecting sets were created using the UpSetR package (Lex *et al*. 2014). Candidate genes within a 20kb interval centered on the peak SNP with p-values above a Benjamini-Hochberg false discovery rate (FDR) threshold of 0.1 were examined further.

#### Comparison to published genome-wide associations in common bean

Out of the 21 BLUPs estimated from CDBN phenotypes, a group of 13 also had published associations from GWAS on common bean. To compare the major associations in our study to those of published studies on balanced genetic trials, we collected the major associations reported in eleven published GWAS studies of common bean (Cichy *et al*. 2015; Kamfwa *et al*. 2015b; Kamfwa *et al*. 2015a; Moghaddam *et al*. 2016; Soltani *et al*. 2017; Tock *et al*. 2017; Nascimento *et al*. 2018; Soltani *et al*. 2018; Oladzad *et al*. 2019a; Oladzad *et al*. 2019b; Raggi *et al*. 2019). We compared these published associations to the associations for the top 10 SNPs for each of the 13 phenotypes in this study, thinned to one SNP per 20kb region. Unfortunately, these comparisons were likely very conservative, in that most of these publications used panels of common bean that were comprised of material from different genepools than the CDBN, with the exception of the MDP and DDP (Moghaddam *et al*. 2016; Soltani *et al*. 2016; Oladzad *et al*. 2019a; Oladzad *et al*. 2019b). Both Andean and Middle-American genepools have been observed to have different SNPs underlying domestication traits (Schmutz *et al*. 2014). Eight of these publications used v1.0 of the *Phaseolus vulgaris* genome annotation, while our associations were mapped to v2.0. We used the genome browser located at https://legumeinfo.org/genomes/gbrowse/phavu.G19833.gnm2 to convert associations between these two versions of the genome annotation. We then determined the number of overlapping associations meeting two criteria: first, those within 200kb of one another, and second, within 20kb of one another and with the same candidate gene. We determined these overlaps for the 80 associations from the eleven published GWAS to find an expected rate of overlap, then compared this to the rate of overlap between this study and the eleven balanced GWAS.

#### Analysis of pleiotropy or linked effects on multiple phenotypes

To increase our power to detect associations above a FDR, and to find genomic associations with significant effects on one or more CDBN phenotype, we used a two-step empirical Bayes procedure, mash, to estimate effects of ∼45000 SNPs on 20 BLUPs determined from CDBN phenotypes (Urbut *et al*. 2019). Mash has been used to increase power to detect effects in analyses of human data, and while the methods are extensible to any dataset with many SNPs/markers and many phenotypes/conditions, it has not yet been used in an agricultural setting. Briefly, mash is a flexible, data-driven method that shares information on patterns of effect size and sign in any dataset where effects can be estimated on a condition-by-condition basis for many conditions (here, phenotypes) across many units (here, SNPs). It first learns patterns of covariance between SNPs and phenotypes from SNPs without strong effects, then combines these data-driven covariances with the original condition-by-condition results to produce improved effect estimates. In this way, mash shares information between conditions to increase the power to detect shared patterns of effects. Importantly, this method does not have restrictive assumptions about the patterns of effects between markers or conditions. In addition, estimates with little uncertainty are not adversely affected by the inclusion of estimates with high uncertainty. Thus, we included 20 phenotypes in the mash analysis, including twelve phenotypes with no signal above the Benjamini-Hochberg FDR threshold in individual GWAS. Two low-signal phenotypes related to bean common mosaic virus presence or absence were not included; inclusion of these phenotypes did not significantly alter the mash results (data not shown). The procedure we used to generate input matrices for mash is captured in the R package gapit2mashr, available at https://github.com/Alice-MacQueen/gapit2mashr. Briefly, the effect of the alternate allele relative to the reference allele was determined for each SNP using GAPIT. To allow mash to converge effectively on effect estimates, the effects for each phenotype were standardized to fall between -1 and 1, with a mean of 0. Because mash does not accept NA values, when GAPIT calculated standard errors for 95% or fewer of the SNPs in the GWAS, we instead calculated standard errors for that phenotype using Hedges’ G (Hedges and Olkin 1985).

Data-driven covariance matrices were estimated using 45,000 randomly selected SNPs from the entire set of 1,221,540 SNPs. These matrices were then used on the top 4,000 SNPs for each of the 20 traits, as determined by *p*-value in the individual GWAS, which produced a matrix of strong effects for 45,000 SNPs. We then explored the patterns of significant effects in the mash output. We first determined which SNPs had evidence of significant phenotypic effects by determining SNPs with the largest Bayes factors. In this analysis, the Bayes factor was the ratio of the likelihood of one or more significant phenotypic effects at a SNP to the likelihood that the SNP had only null effects. Here, following Kass and Raftery (1995), a Bayes factor of > 10^2^ is considered decisive evidence in favor of the hypothesis that a SNP has one or more significant phenotypic effect. We also compared the size of significant phenotypic effects, as determined by SNPs with a local false sign rate of 0.05 or less for one or more phenotype.

The local false sign rate is analogous to a FDR, but is more conservative, in that it also reflects the uncertainty in the estimation of the sign of the effect (Stephens 2017).

#### Data availability statement

Genotypic data is available on SRA under submission number SUB6162710. Code for SNP calling is available at https://github.com/Alice-MacQueen/SNP-calling-pipeline-GBS-ApeKI. Aligned SNP data is available at https://doi.org/10.18738/T8/RTBTIR. Raw phenotypic data is available in the National Agricultural Library: https://www.nal.usda.gov/. Code used to generate data used in this analysis from the raw phenotypic data is available at Rpubs, found at: http://rpubs.com/alice_macqueen/CDBN_Phenotype_Standardization. Code and data necessary to replicate this analysis are available as part of the R package CDBNgenomics, found at: https://github.com/Alice-MacQueen/CDBNgenomics. Supplementary data for this manuscript is available at: https://doi.org/10.18738/T8/KZFZ6K.

## Results

### Cooperative Dry Bean Nursery selection framework

Selection and breeding strategies to generate new bean entries for the CDBN varied across years and among breeding programs. However, in general, new advanced lines were selected from either single, triple, or double crosses among advanced breeding material and released cultivars, which in most cases were already tested within the CDBN in previous years. These lines were bulked to increase seed supply, then field tested to ensure consistency of phenotypic responses in the advanced lines. Entries with favorable characteristics were often entered into the CDBN to be phenotyped in multiple environments. Consequently, most CDBN entries are members of a complex pedigree which has had novel, favorable alleles recombined or introgressed into it over time.

It is clear that the CDBN is not a randomly mating, homogeneous population, and the breeding and selection strategy in the CDBN likely impacts GWAS on this material in a number of ways. Presumably, breeders have increased the frequency of alleles that favorably affect phenotypes over time, which should aid in the detection of these genomic regions via GWAS. The multiple generations of inbreeding should reduce allelic heterogeneity, which should also aid GWAS. Indeed, we find few heterozygous regions in our SNP dataset, and few examples of multiallelic loci. By the same token, the frequent inbreeding may also increase the size of linkage disequilibrium (LD) blocks or cause spurious patterns of LD, which may cause non-syntenic associations and make candidate gene identification more difficult. In addition, the infrequent crosses between the gene pools from the two independent domestication events, and the assortative mating practiced as part of the breeding strategy, could lead to an inflated false positive rate and create correlations between previously uncorrelated traits (Li *et al*. 2017).

### Phenotypic Correlations in the Cooperative Dry Bean Nursery

The CDBN contains a wealth of data to study the genetics of phenotypes and phenotypic correlations (Fig. 1, Fig. 2a). We were able to obtain and genotype 327 germplasm entries from the 544+ entries present in the CDBN trials from 1981 to 2015, including 124 entries that were neither released commercially nor submitted to the National Plant Germplasm System (Npgs 2017), and 39 entries whose seed would not germinate. Most of the remaining entries were grown in the CDBN before 1990 and had seed stocks that, for reasons of practicality, were no longer maintained by breeders (Fig. 1). GBS of the available genotypes generated 1.2M SNPs for analysis of stable effects in the CDBN.

BLUPs of phenotypes from the CDBN, conditioned on location, location by year, and the kinship matrix, are analogous to breeding values for the CDBN entries. These genetic values can be used to determine the narrow-sense heritability, h^2^, potentially explainable by GWAS. h^2^ varied between 6% and 73% in the 21 phenotypic BLUPs (Table 1). We then determined the correlations between the BLUPs of CDBN phenotypes, or the genetic correlations. Correlation coefficients between BLUPs of CDBN phenotypes varied between -0.75 and 0.81, and most phenotypes were significantly correlated (Figure S1). Two major groups of phenotypes were positively correlated: biomass, days to flowering, plant height, zinc deficiency score, days to maturity, blackroot presence/absence, and early vigor were in the first of these groups, and white mold damage score, growth habit, seed yield, harvest index, lodging, rust damage score, bean common mosaic virus damage score, and halo blight damage score were in the second of these groups. These two groups had negative phenotypic correlations with each other.

**Table 1.** Best linear unbiased predictor statistics, with phenotypes ordered as in Figure 2.

| Phenotype | $V_g$ | $V_e$ | $h^2$ | BLUP Range | Units |
| --- | --- | --- | --- | --- | --- |
| Seed yield (kg/ha) | 53173 | 222409 | 0.193 | 1727 | kg ha<sup>-1</sup> |
| Seed weight (mg) | 2340 | 1076 | 0.685 | 443 | mg |
| Days to maturity | 12.3 | 18.3 | 0.402 | 17.0 | days |
| Days to flowering | 4.68 | 6.11 | 0.434 | 11.7 | days |
| Seedfill duration (days) | 5.584 | 17.812 | 0.239 | 12.7 | days |
| Lodging score | 0.244 | 0.466 | 0.344 | 2.97 | 1-5 scale |
| Harvest index (%) | 13.5 | 31.5 | 0.299 | 20.1 | % |
| Plant height (cm) | 13.8 | 42.5 | 0.245 | 18.1 | cm |
| Biomass | 2.46E+05 | 6.72E+05 | 0.268 | 2990 | kg |
| Growth habit | 0.160 | 0.154 | 0.509 | 2.19 | 1-3 scale |
| Seed appearance score | 0.017 | 0.241 | 0.067 | 0.346 | 1-3 scale |
| CBB damage score | 0.282 | 1.127 | 0.200 | 2.32 | 1-9 scale |
| Rust damage score | 3.201 | 1.957 | 0.621 | 7.35 | 1-9 scale |
| Early vigor score | 0.064 | 0.747 | 0.078 | 0.957 | 1-9 scale |
| White mold damage score | 0.143 | 0.631 | 0.185 | 2.60 | 1-5 scale |
| CTV presence/absence | 0.025 | 0.132 | 0.157 | 0.432 | 0-1 scale |
| Halo blight damage score | 0.176 | 0.708 | 0.199 | 1.12 | 1-5 scale |
| BCMV blackroot response | 0.049 | 0.086 | 0.363 | 0.843 | 0-1 scale |
| BCMV presence/absence | 0.016 | 0.097 | 0.145 | 0.428 | 0-1 scale |
| Root rot damage score | 0.455 | 3.193 | 0.125 | 2.25 | 1-9 scale |
| Zinc deficiency damage score | 3.831 | 1.395 | 0.733 | 8.96 | 1-9 scale |

$V_g$ is the REML estimate of the genetic variance from rrBLUP. $V_e$ is the REML estimate of the
error variance. $h^2$ is narrow sense heritability, defined as $V_g / (V_g + V_e)$ .

### Eight CDBN phenotypes have genetic associations above the false discovery rate

We conducted GWAS on 21 phenotypes using best linear unbiased predictors (BLUPs) calculated using a kinship matrix, location, and an interaction between location and year as fixed effects. (for details, see the *Genome-wide association study* section in the Materials and Methods). To determine if any SNP frequencies had changed over the duration of the CDBN, we also conducted GWAS on the earliest year that each germplasm entry was present in the CDBN as a proxy for the age of the entry. This GWAS was analogous to an environmental GWAS that uses climatic variables associated with a genotype’s location of origin (Hancock *et al*. 2011; Morris *et al*. 2013), though this GWAS is fitted to a variable correlated with the age of the genotype rather than with its location of origin.

Given the analytical issues surrounding the use of METs for unplanned genetic analyses, it was unclear whether GWAS on CDBN phenotypes would find significant associations, or if these associations would be reduced or eliminated by environmental noise or by experimental design biases. Thus, we determined if any GWAS on CDBN phenotypes had significant associations after a Benjamini-Hochberg FDR correction of 10%. With this criterion, significant associations were discovered for eight of the 21 phenotypes. More than 33 peaks had SNPs with *p*-values above the FDR, indicating the presence of 30 or more distinct, significant associations with these eight CDBN-derived phenotypes. Phenotypes with associations above the FDR generally had more datapoints in the CDBN (6500 vs 2400 datapoints, Wilcoxon rank sum test *p* = 0.018; Fig. 2a). Phenotypes with associations above the FDR also had significantly higher narrow-sense heritabilities estimated from the phenotypic data (h^2^ of 40.5% vs 25%, Wilcoxon rank sum test *p* = 0.038, Table 1). We briefly discuss the associations above the FDR for these eight phenotypes in the order of most to fewest datapoints in the CDBN. In cases where there were multiple associations for a single phenotype, we discuss only the top associations by *p*-value.

Seed yield (kg ha^-1^) had one significant peak after FDR correction, on Pv01 at 42.2Mb (Fig. 2b, 2c, Table S4). This association was correlated with a difference in seed yield of 104 kg ha^-1^ (Fig. 2f, Supplementary Table 4). Median seed yield in the CDBN for the Durango, Mesoamerican, and Nueva Granada races was 2803, 2443, and 2038 kg ha^-1^, respectively; thus, this genomic region accounts for changes in seed yield of 3.7-5.1%, or three to four years of improvement effort at historical rates of bean improvement (Vandemark *et al*. 2014). This association was 3.7kb upstream of the gene *Phvul.001G167200*, a gene that is highly expressed in the shoot and root tips of common bean at the 2^nd^ trifoliate stage of development (O’rourke *et al*. 2014; Dash *et al*. 2016). The *A. thaliana* homolog of this gene, *VERNALIZATION INDEPENDENCE 5* (*VIP5*), affects flowering time by activating Flower Locus C (FLC), which is a repressor of flowering (Oh *et al*. 2004).

Seed weight (mg) had associations on nine chromosomes that were significant after FDR (Fig. 2d, 2e); the strongest of these were on Pv02 (Fig. 2g), Pv03, Pv05, and Pv08, though each explained only 1-2% of the variation in seed weight (Table S4). Because seed weight correlates strongly with population structure in the three bean races and two bean gene pools, seven principal components were used to correct for population structure in this GWAS (Table S3). The association on Pv02 was 5kb upstream of gene model *Phvul.002G150600*, a *Sel1* repeat protein. *Sel1-like* repeat proteins are frequently involved in signal transduction pathways and in the assembly of macromolecular complexes (Mittl and Schneider-brachert 2007). The association on Pv03 was 10kb upstream of gene model *Phvul.003G039900*, a jasmonic acid carboxyl methyltransferase. The association on Pv05 was not within 20kb of any gene. The association on Pv08 fell in the coding sequence of *Phvul.008G290600*, a choline-phosphate cytidylyltransferase highly expressed in many tissues, including roots and pods and seeds at the heart stage and stage 2, or seeds 3 – 4 and 8 – 10mm wide (O’rourke *et al*. 2014; Dash *et al*. 2016).

Days to flowering had one significant peak after FDR, on Pv01 between 13.4 and 17.1 Mb (Figure S2a, Table S4). It was correlated with a difference in flowering time of 2 to 3 days, depending on the population (Figure S3a). A candidate gene model hypothesized to affect days to flowering, *Phvul.001G087500*, is located at 13.76 Mb in the V2.0 annotation for *P. vulgaris*. Gene model *Phvul.001G087500* is an ortholog of *KNUCKLES* (*KNU*), a protein which is part of the *Polycomb repressive complex 2*, a complex that affects both flowering time and floral meristem development (De LUCAS *et al*. 2016). *KNU* is activated in the transition to determinate floral meristem development and functions in a feedback loop that promotes determinate development (Payne *et al*. 2004; Sun *et al*. 2014).

Lodging score, where higher scores indicated more stem breakage near ground level, had associations on three chromosomes that were significant after FDR; one on Pv04 at 2.8 Mb, one on Pv05 at 0.4 Mb, and one on Pv07 at 34.5 Mb (Figure S2b, Table S4). In total, these three associations explained 8% of the variation in lodging (Supplementary Figure 4). The signal on Pv04 fell within gene model *Phvul.004G025600*; the *A. thaliana* homolog of this gene is involved in the biosynthesis of inositol pyrophosphate, a cellular signaling molecule involved in metabolism and energy sensing (Desai *et al*. 2014). The signal on Pv05 fell within gene model *Phvul.005G005400*, a uridine diphosphate glycosyltransferase superfamily protein (Dash *et al*. 2016). The strongest signal for lodging, explaining 3% of the variation, fell in the promoter region of gene model *Phvul*.*007G221800*, which is orthologous to *SUPPRESSOR OF AUXIN RESISTANCE 1* (SAR1). In *Arabidopsis thaliana*, *SAR1* increases plant height and internode distance and appears to affect stem thickness (Cernac *et al*. 1997; Parry *et al*. 2006).

Harvest index, or the ratio of seed yield weight to total above ground biomass, had one significant association on Pv03 at 2.1 Mb (Figure S2c, Table S4). The alternate allele was associated with an increase in harvest index of 1.5 – 3.5%, and associated to bean race (Figure S3b). This allele was 20 kb from gene model *Phvul.003G023000*, a cellulose synthase-like protein highly expressed in green mature pods, whole roots, and leaf tissue at the 2^nd^ trifoliate leaf stage of development (O’rourke *et al*. 2014; Dash *et al*. 2016).

Growth habit encompasses both determinate and indeterminate types (I and II/III), as well as upright and prostrate indeterminate types (II and III). Growth habit had significant associations on every chromosome after FDR; the strongest four associations were on Pv01 at 6.2 and 42.2 Mb, on Pv09 at 30.9 Mb, and Pv10 at 42.7 Mb (Figure S2d, Table S4). There are known to be multiple determinacy loci segregating in different gene pools of common bean (Kwak *et al*. 2012), which could complicate associations between growth habit and genomic regions in the CDBN panel. These four associations were associated with variation in determinacy in this panel; however, these four associations were not sufficient to explain all variation in determinacy, in that 13 genotypes had all alleles that were associated with determinacy, but were indeterminate, and one genotype had all alleles that were associated with indeterminacy, but was determinate (Figure S3c). The association at 6 Mb on Pv01 fell in the coding sequence of the gene model *Phvul.001G055600*, a RING-CH type zinc finger protein expressed highly in roots and in stem internodes above the cotyledon at the 2^nd^ trifoliate stage (O’rourke *et al*. 2014; Dash *et al*. 2016). The association at 42.2 Mb was 3.7 kb upstream of the gene VIP5; as noted above, this gene and genomic region were also candidate associations for seed yield (kg ha^-1^). The association on Pv09 was 5 kb upstream of model *Phvul.009G204100* that encodes a signal peptide peptidase A highly expressed in pods associated with stage 2 seeds and in stem internodes above the cotyledon at the 2^nd^ trifoliate stage (O’rourke *et al*. 2014; Dash *et al*. 2016). The association on Pv10 was 1 kb upstream of model *Phvul.010G146500*, a gene from an uncharacterized protein family highly expressed in roots, pods with seeds at the heart stage, and stem internodes above the cotyledon at the 2^nd^ trifoliate stage (O’rourke *et al*. 2014; Dash *et al*. 2016).

Bean rust (*Uromyces appendiculatus*) causes leaf and pod pustules and leads to losses in vigor and seed yield. Higher plant damage caused by rust was indicated by a higher rust score. Rust score had significant associations on ten chromosomes after FDR (Figure S2e, Table S4). However, the strongest association was located on Pv11 at 50.6 Mb and overlapped a major cluster of disease resistance genes containing the rust resistance genes *Ur-3*, *Ur-6*, *Ur-7*, and *Ur-11* (Hurtado-gonzales et al. 2017). This signal fell just upstream of the gene model *Phvul.011G193100*, which maps in the interval suggested to contain the resistance gene *Ur-3* (Hurtado-gonzales *et al*. 2017). The alternate allele was present in the early years of our CDBN data within Mesoamerican race, but was either absent or rare within the Durango race in the CDBN until 1988, when it appeared in the pinto Sierra and the great northern Starlight. The alternate allele was not widely distributed in the Durango race until the mid-1990’s (Figure S3d).

Finally, the presence or absence of curly top virus, a virus characterized by plant stunting and deformation of leaves and fruit, had significant associations on seven chromosomes after FDR; however, the strongest associations were on Pv01, Pv05, Pv07, and Pv11 (Figure S2f, Table S4). The association on Pv01 was 0.5 kb upstream of gene model *Phvul.001221100*, recently identified as the photoperiod sensitivity locus *Ppd*, or *PHYTOCHROME A3* (Weller et al. 2019). The association on Pv05 was within 20kb of gene model *Phvul.005G051400*, a VQ motif-containing protein highly expressed in leaf tissue. VQ motif-containing proteins are a class of plant-specific transcriptional regulators that regulate photomorphogenesis and responses to biotic and abiotic stresses (Jing and Lin 2015). The association on Pv07 was 1kb upstream of gene model *Phvul.007G035300*, a pH-response regulator protein. The association on Pv11 was 20kb downstream of gene model *Phvul.011G142800*, a terpene synthase gene expressed in young trifoliates, flowers, and young pods (O’rourke *et al*. 2014; Dash *et al*. 2016). Terpenoids are a large class of secondary metabolite which have roles in plant defense against biotic and abiotic stresses (Singh and Sharma 2015).

#### Three CDBN genetic associations overlap genetic associations from balanced genetic field trials

The presence of many associations above the FDR threshold supports using MET data for genetic analyses. However, the assortative mating employed purposefully by breeders of entries in the CDBN could potentially lead to a high rate of false positives (Li *et al*. 2017). Overall, it was unclear whether GWAS using phenotypes derived from sparse MET datasets would yield similar genetic associations as published, balanced field trials. Thus, we compared the top associations discovered here to associations from eleven published GWAS papers on common bean. This allowed us to compare association overlaps for 13 phenotypes, seven of which that had associations above the FDR, and gave 34 top associations from this study to compare to 80 published association regions. In addition to these GWAS associations, the bean rust resistance phenotype overlapped with a candidate rust resistance gene, *Ur-3*, one of the two genes pyramided early on in bean breeding to provide comprehensive rust resistance.

Three major associations from this study were within 20kb of, and had the same candidate gene as, top associations from published, balanced GWAS: days to flowering, on Pv01 at 13.7 Mb; growth habit, on Pv01 at 42.2 Mb; and lodging, on Pv07 at 34.2 Mb (Table 2). Interestingly, when considering all 114 associations, each of these three regions had significant effects for three phenotypes: lodging, growth habit, and days to flowering on Pv01 at 13.7Mb; growth habit, seed yield, and biomass on Pv01 at 42.2Mb; and plant height, lodging, and growth habit on Pv07 at 34.2Mb (Table 2). In this study, the top 10 SNPs for harvest index and days to maturity also had the same candidate gene on Pv03 at 36.8 Mb, the gene model *Phvul.003G153100*. *Phvul.003G153100* is an AP2-like ethylene-responsive transcription factor highly expressed in root tissue and nodules (O’rourke *et al*. 2014; Dash *et al*. 2016).

**Table 2.**
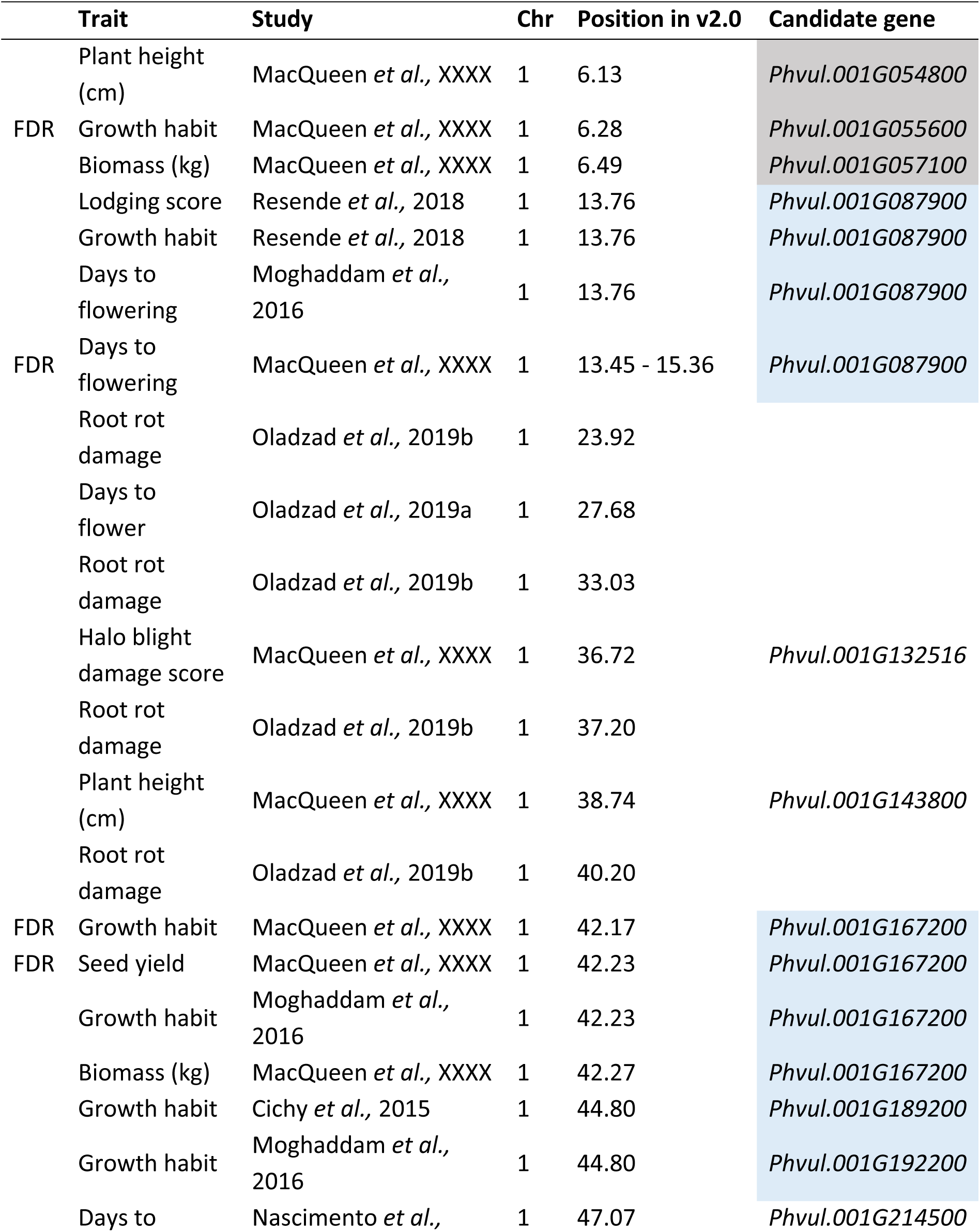
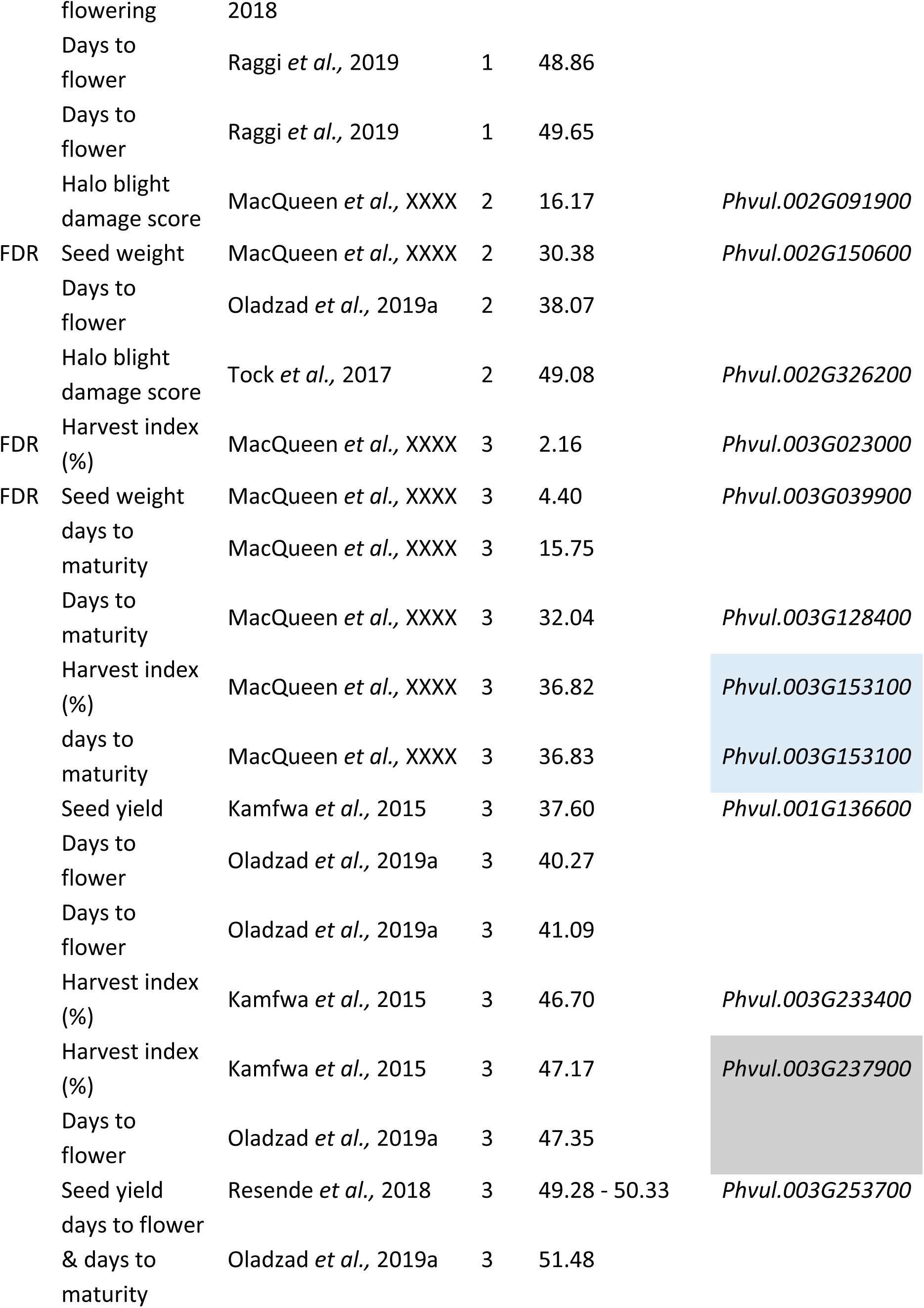
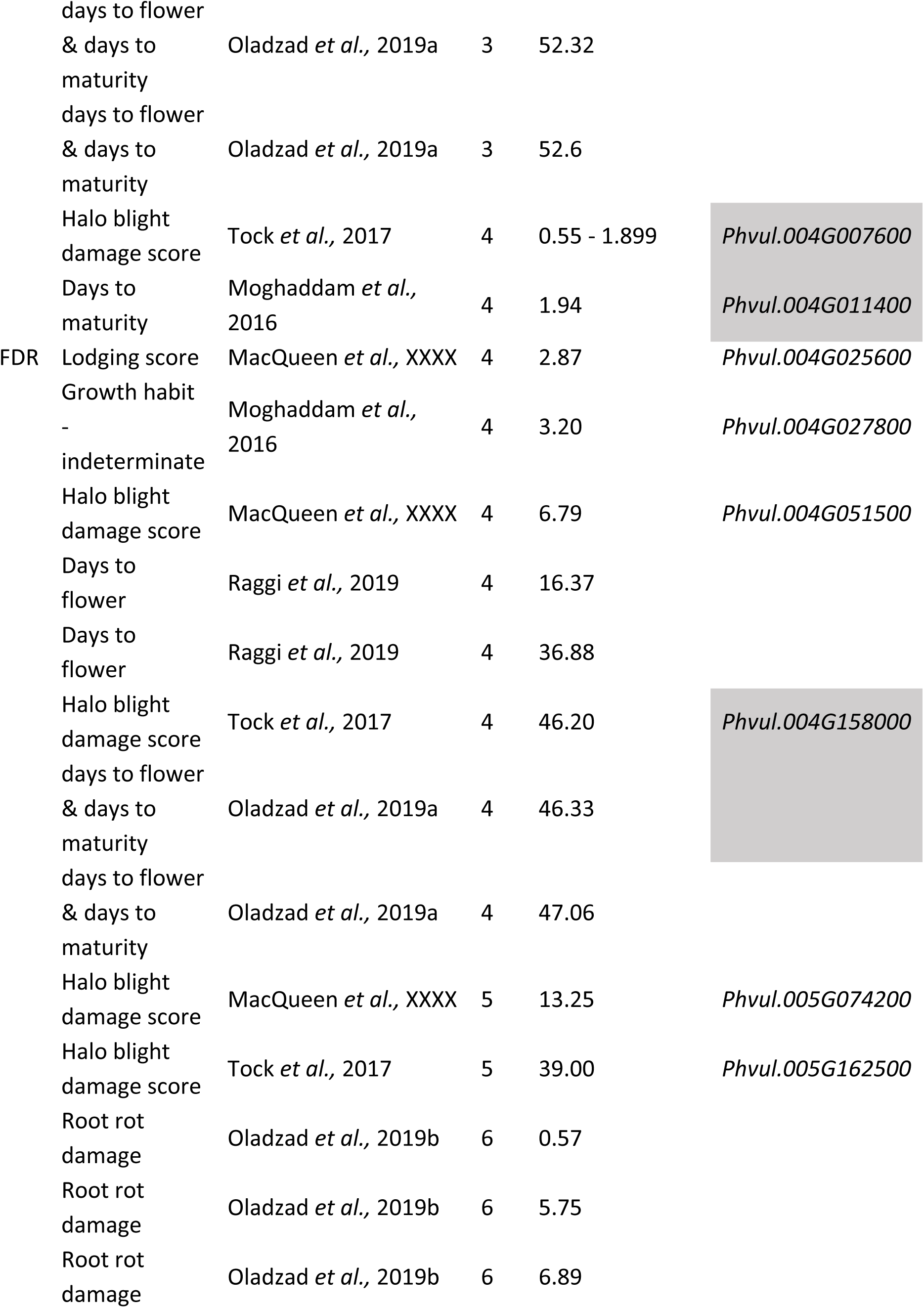
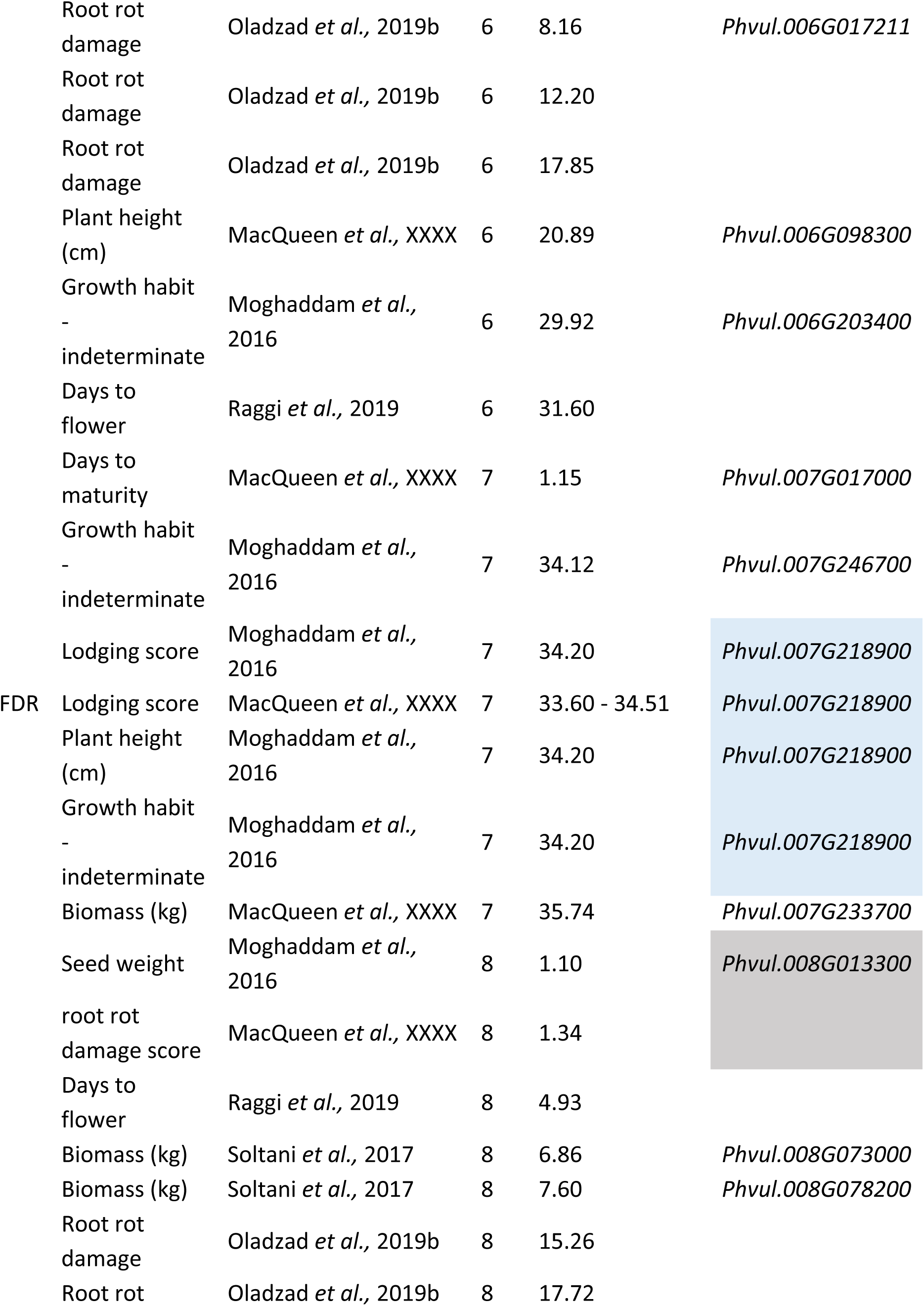
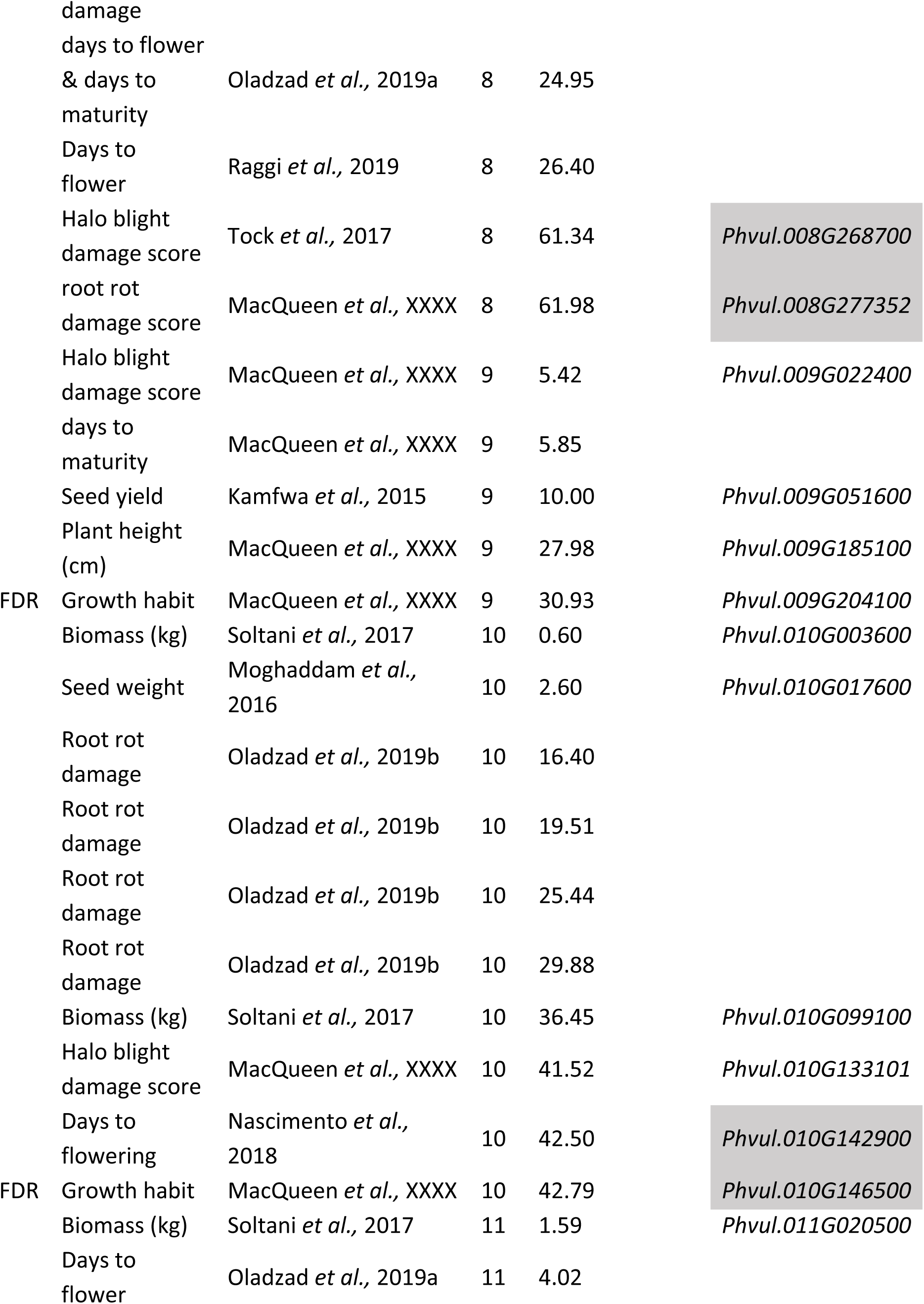
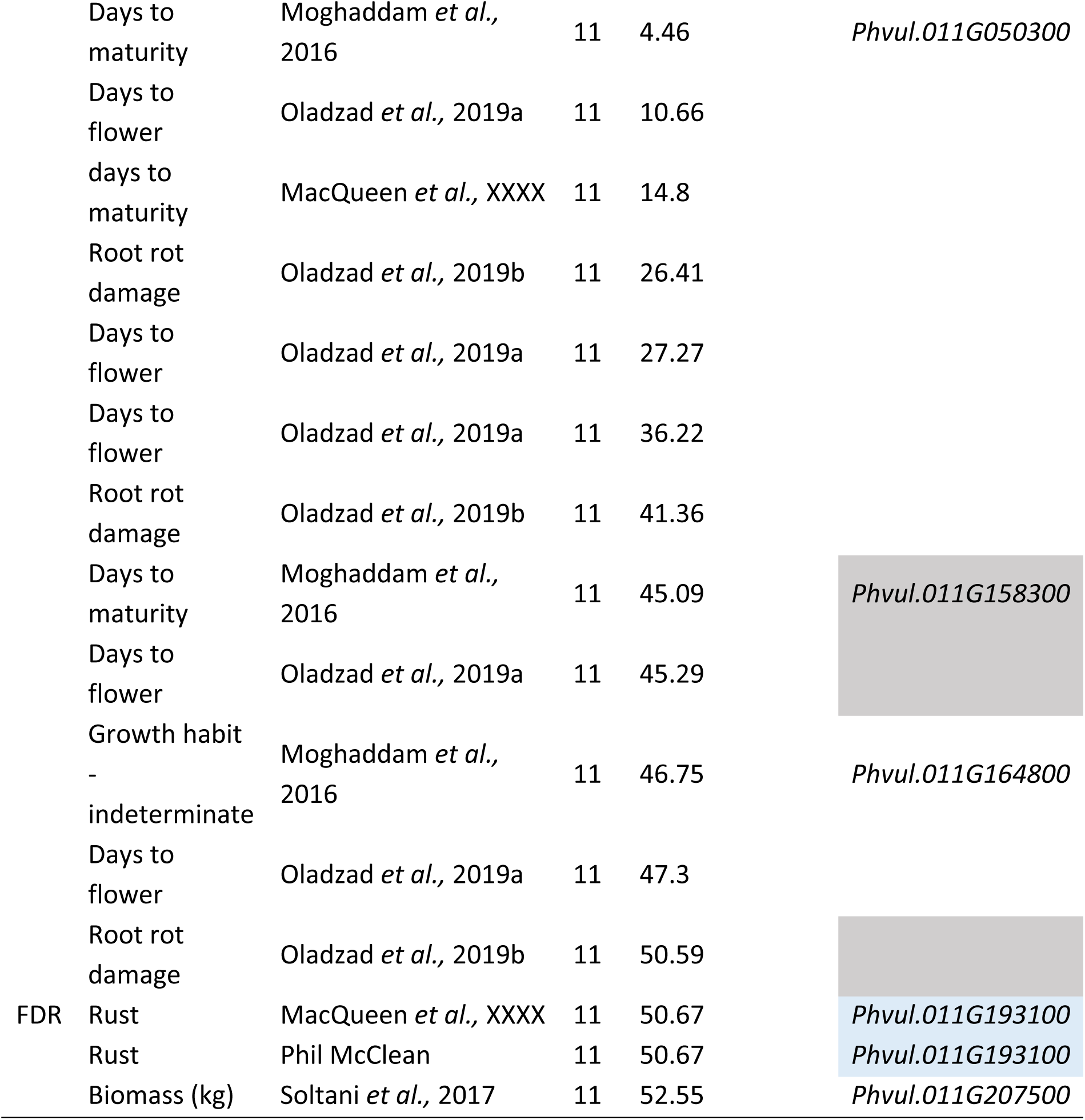
Major associations in genome-wide associations (GWAS) from phenotypes from the Cooperative Dry Bean Nursery (CDBN) and from previously published GWAS. FDR indicates associations from this paper which were above the Benjamini-Hochberg false discovery rate correction. Colors indicate associations in more than one published GWAS: blue indicates associations within 20kb, with the same candidate gene, and grey indicates associations within 200kb.

In comparisons involving only the eleven balanced studies, nine of 80 associations fell into three 20kb regions, while 15 of the 80 associations fell into six 200kb regions. When this study was added, seven additional associations fell into four 20kb regions, while twelve additional associations fell into 14 overlapping 200kb regions (Table 2). This study did not identify many new overlaps at the 20kb level, though it did find associations in all three 20kb overlapping regions found by comparing the eleven balanced studies alone. It did, however, find many new overlaps with previously published studies at the 200kb level, twice as many as expected given the rate of overlap in the eleven balanced studies (chi-squared *p* = 0.025). However, as the balanced studies often did not conduct GWAS on similar phenotypes, our “expected” rate of overlap is likely to be biased. Thus, we consider the fact that this study found the same three 20kb regions that overlap in balanced GWAS comparisons to be stronger evidence than the large number of overlaps at the 200kb level that this panel can yield similar associations to balanced GWAS of common bean diversity panels.

#### Extensive pleiotropy or linked effects within CDBN genetic associations

We observed that numerous CDBN phenotypes had overlapping distributions of significantly associated SNPs. These overlaps could be due to pleiotropy – one genetic locus affecting multiple phenotypes – or due to multiple tightly linked genetic loci affecting multiple phenotypes. To formally compare these overlaps, we used mash on 19 sets of 4,000 SNPs with the smallest *p*-values for phenotypes from the CDBN as well as 4,000 SNPs for the earliest year an entry was grown in the CDBN (Figure 3). Mash shares information about effect sizes of SNPs across all phenotypes, while accounting for data-driven covariances in the patterns of effects (Urbut *et al*. 2019). In contrast to phenotype-by-phenotype analyses, where only eight phenotypes had associations above the FDR, in mash, all twenty phenotypes had SNPs with p-values below the local false sign rate, an analog for the FDR. In addition, SNPs typically had local false sign rates below this threshold for 11-14 phenotypes; thus, there was either extensive pleiotropy or frequent linked effects on multiple phenotypes within entries in the CDBN. SNPs with Bayes factors above ∼10^2^, indicative of decisive evidence favoring that SNP having a significant effect on one or more phenotypes, were distributed very unevenly across the genome, with the vast majority of SNPs clustering within two large regions on Pv01 (Fig. 3b, Table S5). Interestingly, the two largest Bayes factors across all 20 phenotypes were within these two regions, on Pv01 at positions 15.4 Mb and 42.2 Mb. These associations were two that overlapped with top associations from published, balanced GWAS (Table 2). Outside of chromosome Pv01, the most significant Bayes factor was found for a SNP on Pv07 at 14.5 Mb. This SNP was not within 100 kb of any annotated gene.

**Figure 3.**
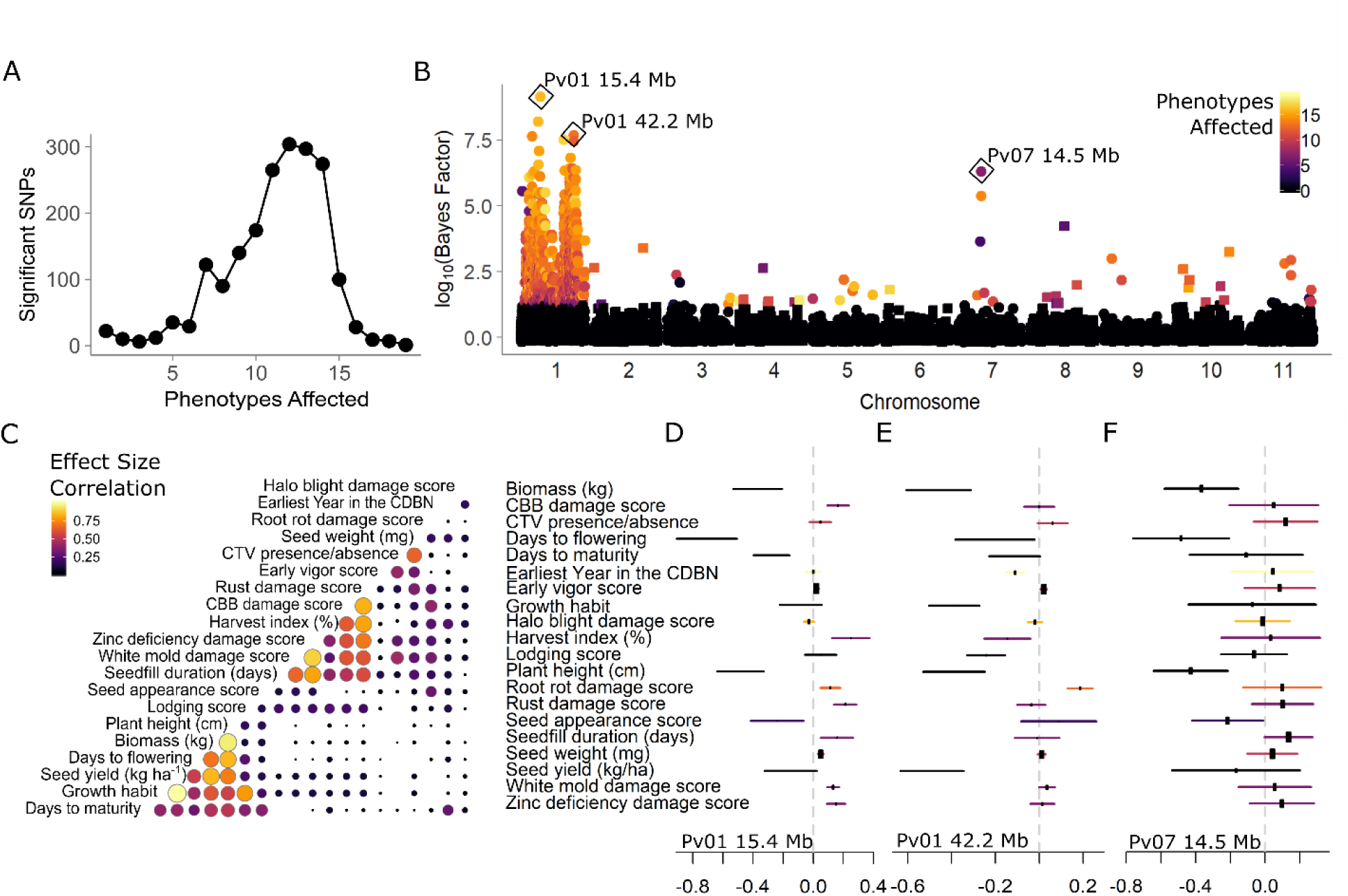
Patterns of phenotypic effects of genetic associations for 22 phenotypes from the Cooperative Dry Bean Nursery (CDBN), determined using multivariate adaptive shrinkage (mash). A) Single-nucleotide polymorphisms (SNPs) with significant effects on one or more of the 22 phenotypes in the CDBN. B) Manhattan plot of the Bayes factor (log_10_) comparing the model likelihood that the SNP has significant effects to the likelihood that it has no significant effects. Bayes factors of > 10^2^ are considered decisive evidence in favor of the alternate model. Point color represents the number of phenotypes for which the SNP has a local false sign rate < 0.05. Squares represent even chromosomes, while circles represent odd chromosomes. The top associations for three regions of the genome are highlighted. C) Correlation in the sign and magnitude of significant effects in all pairwise comparisons of the 22 CDBN phenotypes. Circle size and color indicate the fraction of all significant SNPs that have the same effect sign and similar effect magnitude. D-F) Effect estimates and standard errors for 22 phenotypes for the top associations from three regions of the genome, D) *Phaseolus vulgaris* chromosome 1 (Pv01) at 15.4 Mb, E) Pv01 at 42.2 Mb, F) Pv07 at 14.5 Mb. Genomic locations are based on the *Phaseolus vulgaris* v2.1 genome annotation. Point estimates with higher certainty are indicated by larger rectangles, while standard error bars are colored by the six groups present in C.

The alternate allele for the SNP on Pv01 at 15.4 Mb was associated with significant decreases in biomass, days to flowering, days to maturity, plant height, and seed appearance score. It was also associated with increases in CBB damage score, harvest index, root rot damage score, rust damage score, seed fill duration, white mold damage score, and zinc deficiency damage score (Figure 3d). Here, higher damage scores indicate increased levels of damage. The alternate allele for the SNP on Pv01 at 42.2 Mb was associated with significant decreases in biomass, days to flowering, growth habit (as an increased tendency towards determinacy), harvest index, lodging score, plant height, and seed yield, and increases in root rot damage score (Figure 3e). The allele was also significantly associated with earlier ‘earliest year in the CDBN’, indicating that this allele has been declining in frequency in entries in the CDBN over time. The alternate allele for the SNP on Pv07 at 14.5 Mb was associated with significant decreases in biomass, days to flowering, plant height, and seed appearance score (Figure 3f). Overall, two groups of phenotypes had consistent patterns of effect sign and effect magnitude for most significant SNPs (Fig. 3c). Days to maturity, growth habit, seed yield, days to flowering, biomass, and plant height had a large fraction of SNPs with significant effects with similar effects on these phenotypes; in most pairwise comparisons of these six traits, 40 – 90% of SNPs had the same sign and similar magnitudes of effect (Fig. 3c). The same was true for seed fill duration, white mold damage score, zinc deficiency damage score, harvest index, CBB damage score, and rust damage score; in pairwise comparisons of these six traits, 25 – 80% of SNPs had the same sign and similar magnitudes of effect (Fig. 3c). The phenotypes in the first group corresponded to plant architecture and size, while several phenotypes in the second group were related to disease response. Few other SNPs (∼<10%) affected these two clusters of phenotypes in a similar magnitude with the same sign. Interestingly, groups of highly positively correlated phenotypic BLUPs, or genetic values, did not consistently match groups with large fractions of SNP effects of the same sign and similar magnitude (Figure S4). 90% SNPs with Bayes factors above 10^2^ affected 10 or more phenotypes (Table S5), and typically affected phenotypes in the two groups in similar ways; however, a few exceptions included Pv03 at 10.64 Mb, which affected only plant height; Pv04 at 17.77 Mb, which affected seed weight and varied with earliest year in the CDBN; Pv07 at 13.94 Mb, which affected biomass; and Pv08 at 33.18 Mb, which affected days to flowering, plant height, and seed appearance.

## Discussion

The genes and genomic regions affecting phenotypic variation in common bean are now being narrowed down with the aid of a recently released high-quality reference genome (Schmutz *et al*. 2014). Using previously generated phenotypic data for genetic analysis could circumvent the “phenotypic bottleneck” that has previously constrained our understanding of the genotype-phenotype map in this species. The CDBN offers a vast phenotypic data resource for common bean; however, it was unclear whether the sparse phenotypic data matrix from the CDBN, where only 20 to 30 entries were tested in each location and year, could be used for GWAS. Our results provide evidence supporting the use of METs such as the CDBN for genetic analysis. First, eight of the 22 phenotypes created using the CDBN data had associations that fell above the Bonferroni-Hochberg FDR threshold, and five of these phenotypes had multiple independent peaks that fell above this threshold. Given our FDR of 10%, there were at least 30 distinct, significant associations with these CDBN-derived BLUPs for phenotypes, and these associations tended to be found in phenotypes with higher narrow-sense heritabilities. However, it is still surprising that only eight of the 22 phenotypes had significant associations by the FDR criterion.

We hypothesized that noise caused by environmental variation in phenotypes across years and locations reduced our ability to find significant associations in a condition-by-condition analysis. Supporting this hypothesis, we found that phenotypes with more datapoints in the CDBN were more likely to have associations above the FDR. Thus, we used mash to increase our power to detect significant effects for 20 of these phenotypes, and used an analogue of the FDR, the local false sign rate, to determine whether an effect was significant. By combining information about phenotypic effects across correlated phenotypes, we found significant associations for all phenotypes included in the mash analysis. Thus, phenotypes derived from CDBN MET data are suitable for analysis using GWAS, and the additional phenotypic data available in this MET can be analyzed in mash to boost the power to detect significant genetic effects for traits with pleiotropic genetic architectures.

Second, associations found in our GWAS coincided with results of previous GWAS using balanced phenotypic datasets. Three associations from this study overlapped top associations from published, balanced GWAS: Pv01 at 13.7 Mb, Pv01 at 42.2 Mb, and Pv07 at 34.2 Mb (Table 2). The association at 13.7 Mb fell near the candidate gene *KNU*, a gene which is activated in, and later promotes, the transition to determinate floral meristem development. This peak falls within an association for days to flowering observed previously (Moghaddam *et al*. 2016). The association at 42.2 Mb fell near the candidate gene *VIP5*, an important regulator of flowering time in *A. thaliana* and other species (Huang *et al*. 2012). Other mapping studies have also co-located *VIP5* with QTL for flowering time (Zhou *et al*. 2014). The association at 34.2 Mb on Pv07 also overlapped the strongest association for the earliest year each entry was grown in the CDBN, a proxy for the age of the CDBN entry. This association fell near the candidate gene *SAR1*, which increases plant height and internode distance in *A. thaliana* (Cernac *et al*. 1997; Parry and Estelle 2006). The alternate allele for the signal on Pv07 occurred in newer CDBN entries.

Third, our results are consistent with the recent history of breeding efforts in common beans and provide a map of the genomic regions that have been associated with improvement in the species. We find two major genomic regions on Pv01 associated with many CDBN phenotypes (Figure 3b), which we suggest were major targets of selection by breeders for entries that match an ‘ideotype’ for common bean. The original ideotype had a long hypocotyl, many nodes carrying long pods and without side branches, small leaves, and determinate growth (Adams 1982; Kelly 2001). The primary plant architecture change introduced into genotypes tested in the CDBN over the past 30 years was the adoption of upright indeterminate architecture (Type II), which replaced upright determinate (Type I) architecture in the Mesoamerican race and was introduced into prostrate indeterminate (Type III) germplasm (Kelly 2001; Soltani *et al*. 2016). Generally, entries with Type II architecture yielded more than determinate (Type I) entries, due to the increased pod set associated with indeterminate growth (Kelly 2001), and could yield more than Type III entries under grower-preferred direct harvest (Eckert *et al*. 2011). An association for growth habit on Pv01 at 42.2 Mb fell near the gene *VIP5*; this SNP and gene were also candidate associations for seed yield in this study and days to flowering in Moghaddam *et al*. (2016). The Pv01, Pv09, and Pv10 associations for growth habit, specifically, variation in determinacy, segregate in different genotypes, consistent with the known multiple origins of determinacy segregating in this species (Figure S3c). However, these associations were not sufficient to explain all variation in determinacy present in this panel, perhaps due to the relative rarity of some variants controlling determinacy within the CDBN panel.

Bean breeders in North American generally avoided modifying days to flowering over the years of the CDBN, to protect matching of phenology to specific production environments. However, when Type II architecture was introduced from Mesoamerica race into the Durango/Jalisco race, the first entries with this architecture showed delayed flowering (Vandemark *et al*. 2014). Our strongest association for days to flowering was near the candidate gene *KNU.* This gene is a candidate for the gene *Higher response* (*Hr)* (Gu *et al*. 1998), which affects flowering time. A BLAST analysis of RAPD primers from previous work constrains the location of *Hr* between 1.4 and 21Mb on Pv01 (Gu *et al*. 1998). *Hr* is thus a plausible candidate for the peak at 13Mb. *Hr* is known to be in LD with the common bean gene *terminal flower 1 (PvTFL1* or *fin*) on Pv01, a major determinacy gene in common bean, (Repinski *et al*. 2012). Thus, this gene could plausibly have been introduced during the introduction of Type II architecture.

The primary disease resistance phenotype introduced into entries in the CDBN over the past 30 years was bean rust resistance. Bean rust (*Uromyces appendiculatus*) was a major disease in North America in the 20^th^ century (Zaumeyer 1947). Though the first rust resistant varieties were released in the 1940’s (Zaumeyer 1947), rust was primarily controlled by chemicals prior to the concerted introduction of rust resistance genes in the mid-1980s (Kelly 2001). Our strongest association for rust damage score fell just upstream of the gene model *Phvul.011G193100*, which maps in the interval suggested to contain the resistance gene *Ur-3* (Hurtado-gonzales *et al*. 2017). Initially described by Ballantyne (1978), *Ur-3* was the first gene aggressively used by US breeders to address bean rust in the mid-1980s (Hurtado-gonzales *et al*. 2017). Combining *Ur-3* and *Ur-11* provides resistance against all known rust races (Pastor-Corrales *et al*. 2003), and the two genes formed the basis of breeding efforts to pyramid major bean rust resistance genes that led to the release of pinto, great northern, and black bean germplasm currently used in breeding programs. The alternate allele was present in the early years of the CDBN data in the Mesoamerican race but was either absent or rare in the Durango/Jalisco race in the CDBN until 1988, when it appeared in the pinto Sierra and the great northern variety Starlight. The alternate allele was not widely distributed in the Durango/Jalisco race until the mid-1990’s (Figure S3d). These results agree with the known timing of breeding for rust resistance.

Finally, this work allowed us to characterize the patterns of sharing of genetic effects on phenotypes in the CDBN. Selection for the common bean ideotype is known to have led to pleiotropic effects on, and associations with other traits, such as seed yield, biomass, and plant height (Soltani *et al*. 2016). Previous work indicated that genes responding to photoperiod have a major influence on many traits, including biomass, harvest index, days to maturity, and plant architecture traits such as the number of branches and nodes (Wallace *et al*. 1993; Gu *et al*. 1994). Our associations also revealed substantial overlaps in the genomic regions affecting phenotypic variation, suggesting the presence of substantial pleiotropy or linked genes of major effect. The genomic region on Pv01 from 34 – 48 Mb has also been identified in previous QTL mapping studies as one that affects many traits, including seed yield, days to flowering, days to maturity, seed fill duration, seed weight, biomass, and pod wall ratio (Trapp *et al*. 2015; Trapp *et al*. 2016). Our mash analysis reveals two major groups of phenotypes with commonly shared SNP effects, one corresponding to plant architecture and size, and the other related disease response. Very few SNPs had similar effects on both groups of traits (Figure 3c). This indicates pleiotropy or correlated effects within each group of phenotypes, and unlinked effects or antagonistic pleiotropy between these groups of phenotypes. In addition, the two groups of phenotypes that had similar genetic effects at the SNP level did not substantially overlap groups of phenotypes with highly correlated genetic values by BLUP estimation (Figure S4). Though many genomic regions affect multiple phenotypes in the CDBN, the large shared effects detected by mash do not always combine additively into the overall patterns of genetic correlation present in this dataset. However, two sets of phenotypes did have shared SNP effects and similar patterns of phenotypic correlations: lodging, seed yield, and growth, and biomass, plant height, days to flowering, and days to maturity. We suggest that these seven phenotypes were the most important when breeders selected for preferred common bean ideotypes. In contrast, many of the remaining phenotypes were related to disease damage; these phenotypes might be more affected by epistatic interactions between genomic regions, or by tradeoffs across environments.

Overall, METs such as the CDBN offer a remarkable opportunity to identify candidate genes underlying phenotypic variation and phenotypic plasticity and to identify how artificial selection has affected crop phenotypes through time. We note that the genomic regions found with this approach are likely to have consistent, stable phenotypic effects across a large range of environments. These genomic regions are thus likely to be generally useful to bean breeding. Detailed mapping and cloning of the causative genes in these regions will provide insight into molecular mechanisms that control these critical phenotypes important for high productivity of common bean. In the future, we also believe that it would be of great value to crop breeding and genetics to archive DNA from all material used in breeding programs and MET trials.

Many crops, both in the U.S. and worldwide, have public trials that could be mined in a manner similar to our approach. This work will require collaborative efforts between crop breeders and bioinformaticians to digitize, clean, and analyze phenotypic data from METs and to obtain genetic material from successful and unsuccessful trial entries. Phenotypic and genetic data can be combined using genomic selection approaches, or by GWAS using models that adjust BLUPs for effects of kinship, trial location, and trial year (Rife *et al*. 2018; Sukumaran *et al*. 2018). If effect estimates for genetic markers can be obtained, and some effects are strong, the patterns of significant effects across markers and phenotypes can be determined using a metanalysis approach such as mash (Urbut *et al*. 2019). A broader effort to collectively mine such extensive phenotypic data could identify conserved genetic factors important for improved productivity for many crops in major production regions.

## Acknowledgements

The authors would like to thank An Hang, John Kolar, and Shree Singh for substantial contributions over the history of the CDBN and Melody Yeh for assistance in digitizing and harmonizing the CDBN reports. USDA is an equal opportunity provider and employer. We thank the Joint Genome Institute for pre-publication access and use of the Phaseolus vulgaris V2.1 genome and annotation. This research was partially funded by support from the National Science Foundation, Plant Genomes Research Program, Grant IOS-1612262 to AHM.

## Author contributions

AHM, JMO, JWW, PEM, and TEJ conceived of the analysis. PEM, JMO, PNM, and JM provided seed for varieties in the CDBN. AHM, RL, PEM, and JS sequenced the CDBN. RL and PEM provided sequence data from the ADP and MDP. JWW compiled the phenotypic data from the annual CDBN reports. AHM conducted the analyses with input from JWW, PEM, and TEJ. AHM wrote the manuscript with contributions from all authors.

## Supporting Information

Supplementary data for this manuscript is available at: https://doi.org/10.18738/T8/KZFZ6K.

**Notes.** Further details on the methods used to generate 22 phenotypes from the Cooperative Dry Bean Nursery dataset of common bean (*Phaseolus vulgaris*). Additional materials and methods concerning phenotypic data processing, greenhouse phenotypes, single nucleotide polymorphism imputation and significance and candidate gene identification criteria.

**Table S1** Summary of the Cooperative Dry Bean Nursery Dataset of common bean (*Phaseolus vulgaris*) phenotypes and the subset used in the present analysis.

**Tables S2** Excel File. Location information, genotyped Cooperative Dry Bean Nursery germplasm information, and corrected phenotype medians for 22 phenotypes used for each entry for genome-wide association on common bean (*Phaseolus vulgaris*).

**Table S3** Number of principle components that maximized the Bayesian Information Criterion for model selection in GAPIT, for each set of BLUPs derived from phenotypes in the Cooperative Dry Bean Nursery dataset.

**Table S4** Excel File. Associations from single phenotype genome-wide association significant using a Benjamini-Hochberg false discovery rate threshold of 10%. Separate tabs of the document are associations for separate phenotypes.

**Tables S5** Excel File. Associations from the multivariate shrinkage analysis significant using a local false sign rate threshold of 5%.

**Fig. S1** Correlations between best linear unbiased predictors (BLUPs) for each phenotyped entry in the Cooperative Dry Bean Nursery.

**Fig. S2** Genomic associations for six additional phenotypes with associations above a Benjamini-Hochberg false discovery rate correction.

**Fig. S3** Specific effects of top associations for four phenotypes in the Cooperative Dry Bean Nursery dataset of common bean (*Phaseolus vulgaris*).

**Fig. S4** Overlap between genetic correlation and effect size correlation groups.

